# Comparison of machine learning to deep learning for automated annotation of Gleason patterns in whole mount prostate cancer histology

**DOI:** 10.1101/2022.11.10.516007

**Authors:** Savannah R. Duenweg, Michael Brehler, Samuel A. Bobholz, Allison K. Lowman, Aleksandra Winiarz, Fitzgerald Kyereme, Andrew Nencka, Kenneth A. Iczkowski, Peter S. LaViolette

## Abstract

**Background:** One in eight men will be affected by prostate cancer (PCa) in their lives. While the current clinical standard prognostic marker for PCa is the Gleason score, it is subject to inter-reviewer variability. This study compares two machine learning methods for discriminating between high- and low-grade PCa on histology from 47 PCa patients.

**Methods:** Digitized slides were annotated by a GU fellowship-trained pathologist. High-resolution tiles were extracted from annotated and unlabeled tissue. Glands were segmented and pathomic features were calculated and averaged across each patient. Patients were separated into a training set of 31 patients (Cohort A, n=9345 tiles) and a testing cohort of 16 patients (Cohort B, n=4375 tiles). Tiles from Cohort A were used to train a compact classification ensemble model and a ResNet model to discriminate tumor and were compared to pathologist annotations.

**Results:** The ensemble and ResNet models had overall accuracies of 89% and 88%, respectively. The ResNet model was additionally able to differentiate Gleason patterns on data from Cohort B while the ensemble model was not.

**Conclusions:** Our results suggest that quantitative pathomic features calculated from PCa histology can distinguish regions of cancer; how-ever, texture features captured by deep learning frameworks better differentiate unique Gleason patterns.

## Introduction

Prostate cancer (PCa) is the most diagnosed non-cutaneous cancer in men, affecting an estimated 268,000 men in 2022[1]. Improved prostate cancer screening and therapies have led to a high five-year survival rate and the overall prognosis for PCa is one of the best compared amongst all cancers. Prostate cancer is currently graded using the Gleason grading system, assigning scores corresponding to the two most predominant morphological patterns present. More recently, it has been used to assign patients into one of five Grade Groups (GG) to predict prognosis[2]. Clinically significant cancer (GG ≥ 2, tumor volume ≥ 0.5 mL, or stage ≥ T3) is often treated with radiation therapy and/or radical prostatectomy. Low-grade cancer can often be monitored through annual prostate specific antigen (PSA) testing. Side effects from prostate cancer treatment can include long-term complications such as impotence and impaired urinary function[3], thus early and accurate detection of PCa is necessary to minimize overtreatment while still addressing clinically significant cancer.

Digital pathology is playing an increasingly important role in clinical research, with applications in diagnosis and treatment decision support[4]. Fast acquisition time, management of data, and interpretation of histology has made digital pathology popular and easier for pathologists to manage and share slides. Additionally, artificial intelligence (AI) with digital pathology has created opportunities to incorporate computational algorithms into pathology workflows or for AI-based computer-aided diagnostics[5].

In prostate cancer research, many machine learning applications have been focused on automated Gleason grading. While the Gleason score is currently the gold standard prognostic marker for prostate cancer, the process of assigning grades is a subjective, quantitative metric. Additionally, pathologist-provided annotations for digital pathology studies is not only time consuming, but can result in significant inter-observer variability[6, 7]. The primary focus of these automated Gleason grading methods has been on biopsies or tissue microarrays as opposed to whole-slide images[8–11]. A fast, automated tool for identifying Gleason patterns in prostate histology could allow for rapid annotation and grading, as well as provide important prognostic information such as recurrence probabilities.

In this study, we developed an Automated Tumor Assessment of pRostate cancer hIstology (ATARI) classification model for the Gleason grading of whole-mount prostate histology using quantitative histomorphometric features calculated from digitized prostate cancer slides. The results of this model were validated using ground truth pathologist annotations. In addition, we compared this model to a residual network with 101 layers (ResNet101) for automated Gleason grading[12]. Specifically, we tested the hypothesis that a machine learning model applied to second-order features calculated from digitized histology could discriminate prostate cancer from normal tissue. We also hypothesized that deep learning model would differ in classification accuracy, both in detecting cancer and differentiating Gleason patterns.

## Materials and Methods

### Patient Population and Data Acquisition

Data from 47 prospectively recruited patients (mean age 59 years) with pathologically confirmed prostate cancer were analyzed for this study. This study was conducted according to the guidelines of the Declaration of Helsinki and approved by the Institutional Review Board of the Medical

College of Wisconsin. Written informed consent was obtained from all subjects involved in the study. The data presented in this study are available on request from the corresponding author. The data are not publicly available due to patient privacy concerns. For model development, subjects were split into 2/3 training (n = 31 patients) and 1/3 testing (n = 16 patients) data sets, matched for tumor grade and other clinical characteristics (see **Table 1**).

**Table 1:**
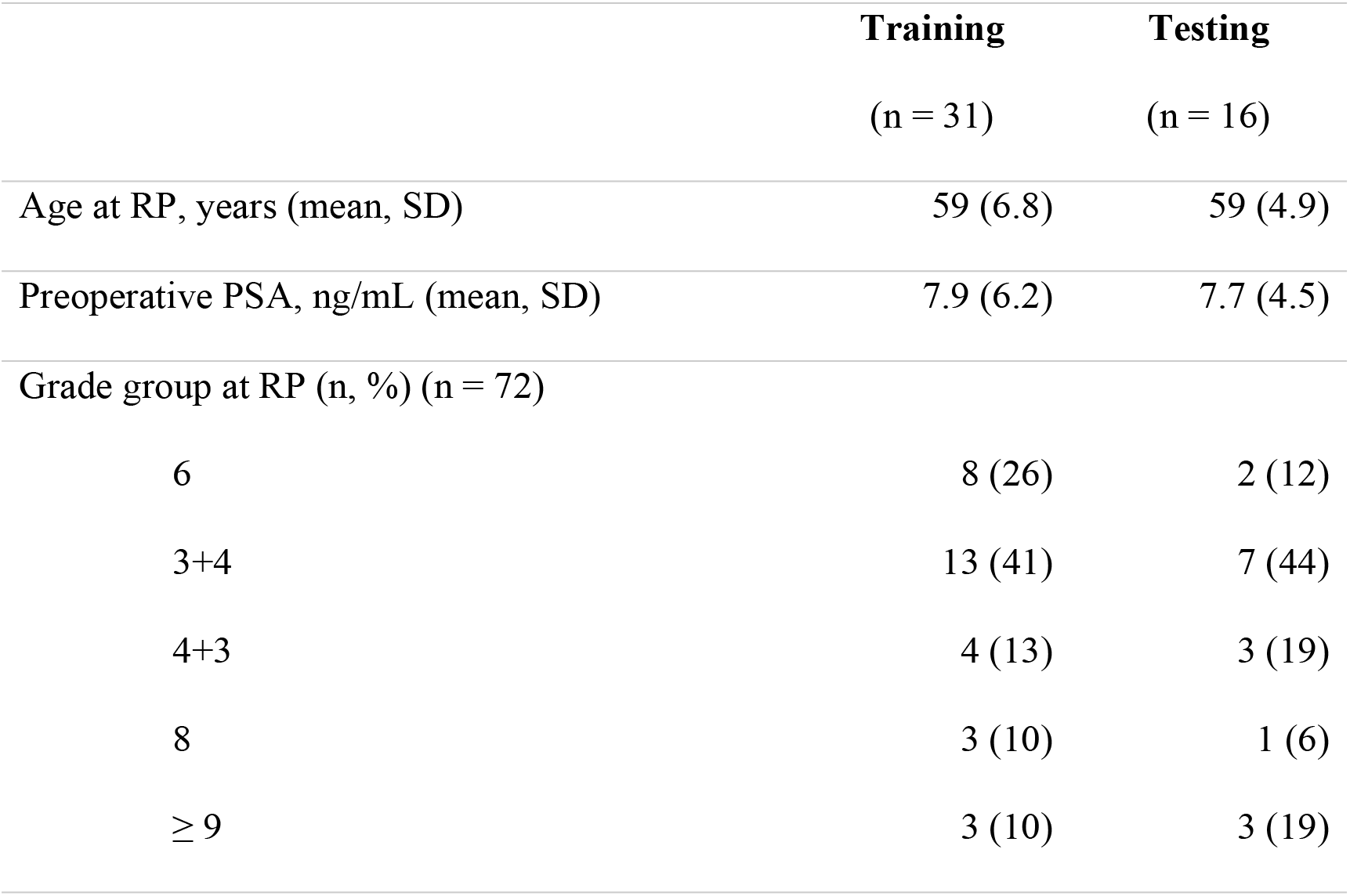
Patient demographics of the study cohort at the time of radical prostatectomy (RP).

|  | **Training** | **Testing** |
| --- | --- | --- |
|  | (n = 31) | (n = 16) |
| Age at RP, years (mean, SD) | 59 (6.8) | 59 (4.9) |
| Preoperative PSA, ng/mL (mean, SD) | 7.9 (6.2) | 7.7 (4.5) |
| Grade group at RP (n, %) (n = 72) |  |  |
| 6 | 8 (26) | 2 (12) |
| 3+4 | 13 (41) | 7 (44) |
| 4+3 | 4 (13) | 3 (19) |
| 8 | 3 (10) | 1 (6) |
| $\geq 9$ | 3 (10) | 3 (19) |

### Tissue Collection and Processing

Prostatectomy was performed using a da Vinci robotic system (Intuitive Surgical, Sunnyvale, CA)[13, 14]. Whole prostate samples were fixed in formalin overnight and sectioned using custom axially oriented slicing jigs[15]. Briefly, prostate masks were manually segmented from the patient’s pre-surgical T2-weighted magnetic resonance image using AFNI (v.19.1.00) (Analysis of Functional NeuroImages, http://afni.nimh.nih.gov/)[16]. Patient-specific slicing jigs were modeled using Blender 2.79b (https://www.blender.org/) to match the orientation and slice thickness of each patient’s T2-weighted image[6, 17–19], and 3D printed using a fifth-generation Makerbot (Makerbot Industries, Brooklyn, NY). The MRI scans were not used beyond slicing molds for the remainder of this study.

Whole-mount tissue sections were processed, paraffin embedded, and resulting whole mount slides were hematoxylin and eosin (H&E) stained. The slides were then digitally scanned using a slide scanner (Olympus America Inc., Center Valley, PA, USA) at a resolution of 0.34 microns per pixel (40x magnification) to produce whole slide images (WSI), and down-sampled by a factor of 8 to decrease processing time. A total of 330 digitized slides were manually annotated using a Microsoft Surface Pro 4 (Microsoft, Seattle, WA, USA) with a pre-loaded color palette for different Gleason patterns[2] by a GU fellowship-trained pathologist (KAI). An example of the prostate annotation process is shown in **Figure 1**.

**Fig 1.**
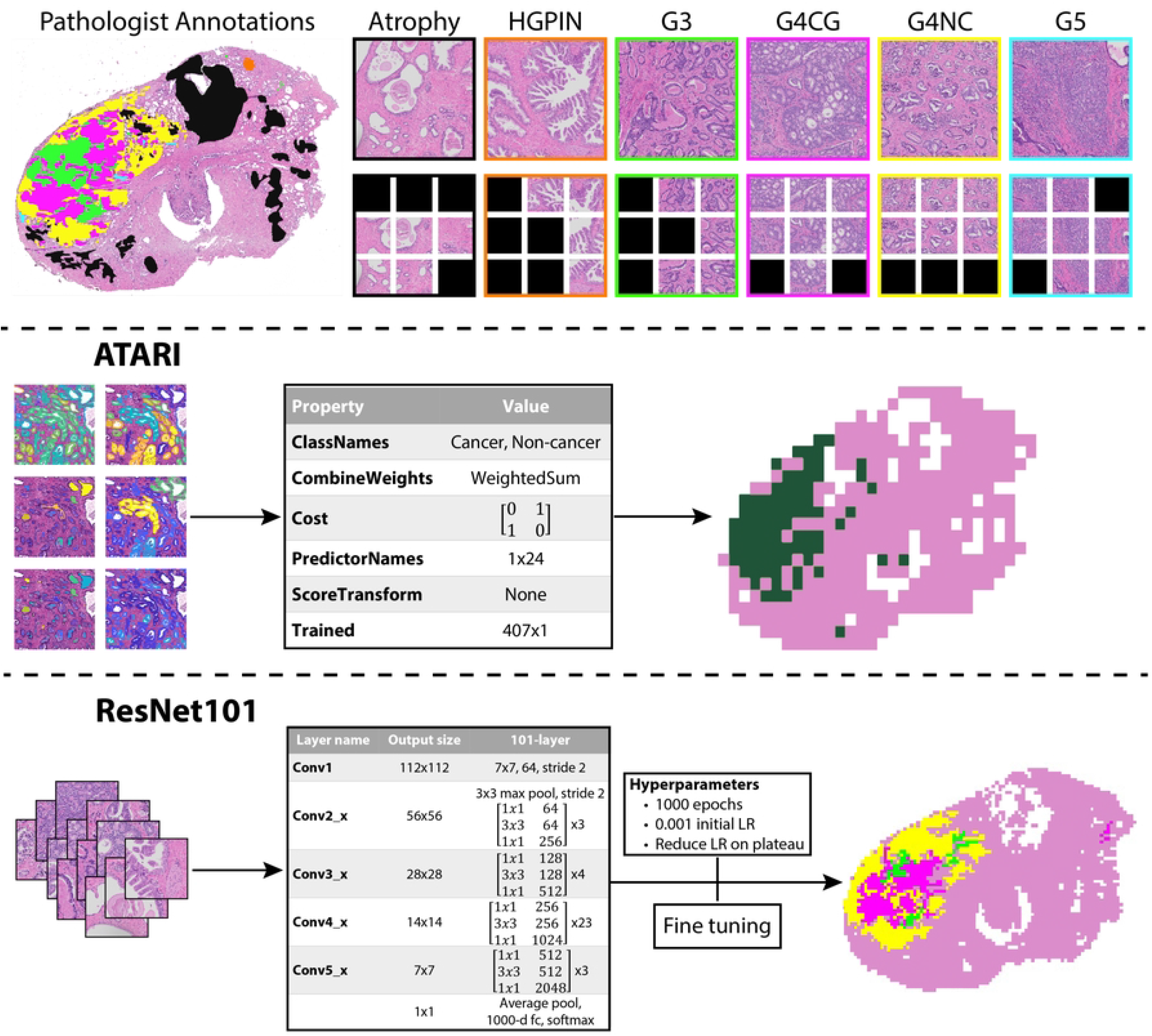
*Top*: Annotation and tile extraction process. After manual annotation of digitized slides, 3000×3000 pixel tiles are extracted from unique annotated regions. Those tiles are then further divided into 1024×1024 pixel tiles and those that remain within a mask are saved (black tiles indicate unsaved tiles). *Middle*: Workflow for the ATARI classifier. Quantitative pathomic features calculated from the large tiles are used as input to a compact classification ensemble to predict cancer vs non-cancer in a whole-slide image. *Bottom*: Workflow for the ResNet101 classifier. 1024×1024 pixel annotated tiles are used as input into the ResNet model to predict non-cancer vs Gleason grade groups. *Abbreviations*: HGPIN = high-grade prostatic intraepithelial neoplasia; G3 = Gleason pattern 3; G4CG = Gleason pattern 4 cribriform; G4NC = Gleason pattern 4 non-cribriform; G5 = Gleason pattern 5.

### Annotation Segmentation

Digital whole-mount slides were divided into high resolution tiles that were 3000×3000 pixels and labeled using their corresponding xy-coordinates within the image. This size tile was chosen as it is the smallest resolution that our pathologist could determine Gleason grades. These tiles were then stitched back together to recreate the whole-mount image while concurrently creating x- and y-coordinate look-up tables. A subset of slides was rescanned on the Olympus slide scanner, and annotations that were performed on lower resolution digitized versions of the slide were quantitatively transferred (n=201 slides). Briefly, the analogous annotated image was aligned to the newly digitized slide using MATLAB 2021b’s *imregister* function (The MathWorks Inc., Natick, MA, USA). The annotations were isolated to create a single mask for each of eight possible classes: seminal vesicles, atrophy, high-grade prostatic intraepithelial neoplasia (HGPIN), Gleason 3 (G3), Gleason 4 cribriform gland (G4CG), Gleason 4 non-cribriform glands (G4NC), Gleason 5 (G5), and unlabeled benign tissue. Gleason 4 patterns have been separated in our annotations as there are notable prognostic differences between the cribriform and non-cribriform patterns[20–23]. An additional averaged white image of non-tissue (i.e., background, lumen, and other artifacts) was found to remove these areas from the annotation masks to ensure the most representative histology remained for analysis. Each region of interest (ROI) within an individual class was individually compared to the xy-look-up tables to determine coordinates corresponding to tiles, and only those with over 50% of a specific annotation were included. Five tiles from each ROI were saved into annotation-specific directories for use with the ATARI model, except for unlabeled benign tissue where 15 tiles were randomly saved from each slide. ROIs that were too small to extract 5 tiles from were excluded.

Each annotated tile was further divided into 1024×1024 pixel tiles for use with the ResNet101 model, resulting in upwards of 9 sub-tiles used for the ResNet101 per full-sized tile used for the ATARI model. Sub-tiles that remained within a mask were saved into annotation-specific directories, similarly to the large tiles used for the ATARI model. The ResNet101 additionally was trained using background tiles determined by areas that were included in the average white image. Tiles used for training were augmented by resizing (250×250 pixel), random cropping (240×240), applying color jitter (0.3, 0.3, 0.3), adding random rotations (±0-30°), applying random horizontal and vertical flips and center cropping to the ResNet input size of 224×224 as well as normalizing to ImageNet’s mean (0.485, 0.456, 0.406) and standard deviation (0.229, 0.224, 0.225). This tile extraction process is demonstrated in **Figure 1**, and breakdown of slides and sorted tiles can be found in **Table 2**.

**Table 2.** Breakdown of tiles used for training and testing each of the models.

|  | **Training**<br>(n = 31) |  | **Testing**<br>(n = 16) |  |
| --- | --- | --- | --- | --- |
| Tissue samples (n, %) | 213 |  | 117 |  |
| Samples per patient (mean, SD) | 6.9 (2.3) |  | 7.3 (1.9) |  |
| Annotated Tiles (n, %) | ATARI | ResNet101 | ATARI | ResNet101 |
| Atrophy | 3555 (38) | 30000 (24) | 1675 (38) | 72098 (57) |
| G3 | 990 (11) | 16000 (13) | 475 (11) | 14565 (11) |
| G4CG | 130 (1) | 5477 (4) | 60 (1) | 1078 (1) |
| G4NC | 515 (6) | 16482 (13) | 235 (5) | 5382 (4) |
| G5 | 75 (1) | 4118 (3) | 55 (1) | 236 (<1) |
| HGPIN | 285 (86) | 4785 (4) | 45 (1) | 610 (<1) |
| Seminal Vesicles | 435 (67) | 10456 (8) | 210 (5) | 5728 (5) |
| Unlabeled Benign Tissue | 3360 (67) | 20000 (16) | 1620 (37) | 13483 (11) |
| Background | 0 (0) | 20000 (16) | 0 (0) | 14027 (11) |
| Total | 9345 | 127319 | 4375 (32) | 127207 |

### Pathomic Feature Extraction

High resolution tiles were down-sampled to increase processing time, and then were processed using a custom, in-house MATLAB function to extract pathological features for use with the ATARI model. First, a color deconvolution algorithm was applied to each image to segment stroma, epithelium, and lumen based on their corresponding stain optical densities (i.e., positive hematoxylin or eosin, and background)[24]. These features were then further smoothed and filtered to remove excess noise and improve segmentations. Glands with lumen touching the edge of a tile were excluded. Overall stromal and epithelial areas were calculated on a whole-image basis, and an additional six features were calculated on an individual gland-basis: epithelial area, roundness, and wall thickness; luminal area and roundness, and cell fraction (i.e., the percent of epithelial cells per total gland area, defined by the area of the epithelium without lumen).

### Model training

Flowcharts for the ATARI model and ResNet101 classifier can be found in **Figure 1**. An ensemble algorithm was used as the framework for developing the ATARI classifier on 31 subjects based in MATLAB (Mathworks, Inc. Natick, MA). A compact classification ensemble was used, which fitted predictors trained on bootstrapped samples from the training data set to obtain a combined ensemble model that minimized variance across learners[25, 26]. Inputs for this model were mean, median, and variance of the calculated pathomic features averaged across each tile, z-scored across the training data. To test the granularity of Gleason pattern prediction, we trained predictive models using several different levels of tumor specificity including all Gleason grades; high-(G4+) and low-grade (G3) cancer and benign tissue (HG vs LG model); and non-cancer and cancer (G3+) (NC vs CA model). To test generalizability, the model was applied to a left-out test set. Predictions were then plotted on three slides from the test data set using the same features calculated across all tiles for the slide to assess successful identification of tumor and compared to ground-truth pathologist annotations and the ResNet model.

To test a deep learning approach for comparison, a ResNet model with 101 layers was implemented in Python using the PyTorch framework (v.1.8.1)[12, 27]. The same tiling procedure as previously described was used to curate the dataset for this network, with the addition of splitting all tiles into smaller 1024×1024 pixel patches and saving those that remained 50% within an annotation mask. Data from Cohort A was split into 80/20 training and validation datasets to prevent overfitting and several data augmentation techniques were used to increase training samples. The image patches were resized to 250×250 pixels, randomly cropped to 240×240 pixels, augmented and center cropped to generate the needed input size of 224×224 pixels. The same three model designs as the ATARI were trained using the ResNet101 framework. Class imbalance of the training dataset was addressed by introducing sample number-based class weights in the cross-entropy loss function.

## Results

The accuracy of both models was analyzed using a left-out test dataset from 17 patients (95,875 image patches for the ResNet; 4,375 image tiles for ATARI). The ATARI model was unable to successfully classify Gleason grades (overall accuracy 85%, per-class accuracy range 0% - 99%) nor high-(HG) and low-grade (LG) cancer (overall accuracy 83%, per-class accuracy range <1% - 99%). In both models, normal tissue was classified well above chance level (20% for all Gleason grades, 33% for high- and low-grade cancer), with G3 in the Gleason grades model and HG in the HG vs LG model performing at chance. The non-cancer (NC) vs cancer (CA) model had an overall accuracy of 89% and a per class accuracy of 97% and 53% for NC and CA, respectively. The ResNet model was able to successfully classify all Gleason grades with an absolute overall accuracy of 79% (per class accuracy range 25% - 87%), HG vs LG (overall accuracy 72%, per class accuracy range 55% - 72%), and NC vs CA (overall accuracy 83%) with an accuracy 91% and 74% for non-cancer and cancer (**Figure 2**).

**Fig 2.**
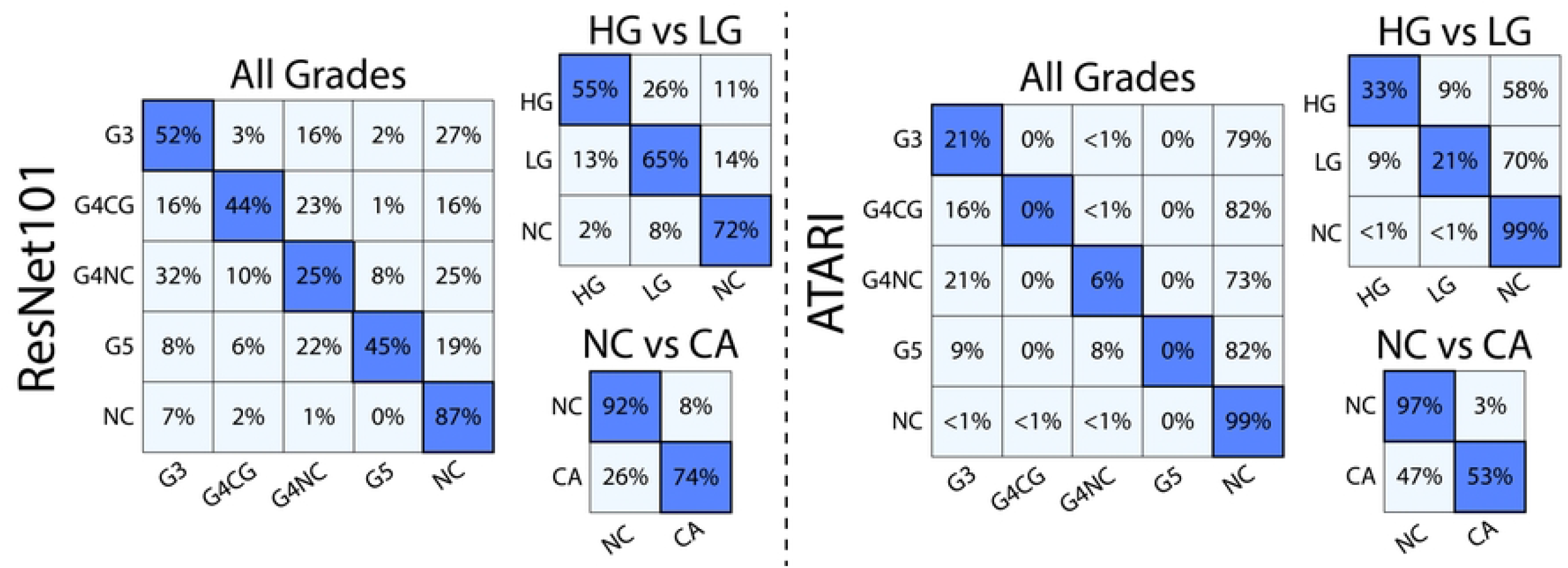
Confusion matrices for the three classification models for both the ResNet101 and ATARI. The ResNet101 was able to distinguish between unique Gleason patterns at higher accuracies that the corresponding ATARI models.

**Figure 3** show the representative slides as their ATARI and ResNet101 annotations as compared to ground-truth annotations. Although the ATARI model was unable to capture unique Gleason patterns, it was able to define the region of tumor present on the slide. The ResNet101 model was able to accurately predict the Gleason patterns with a per class accuracy of 25-52%.

**Fig 3.**
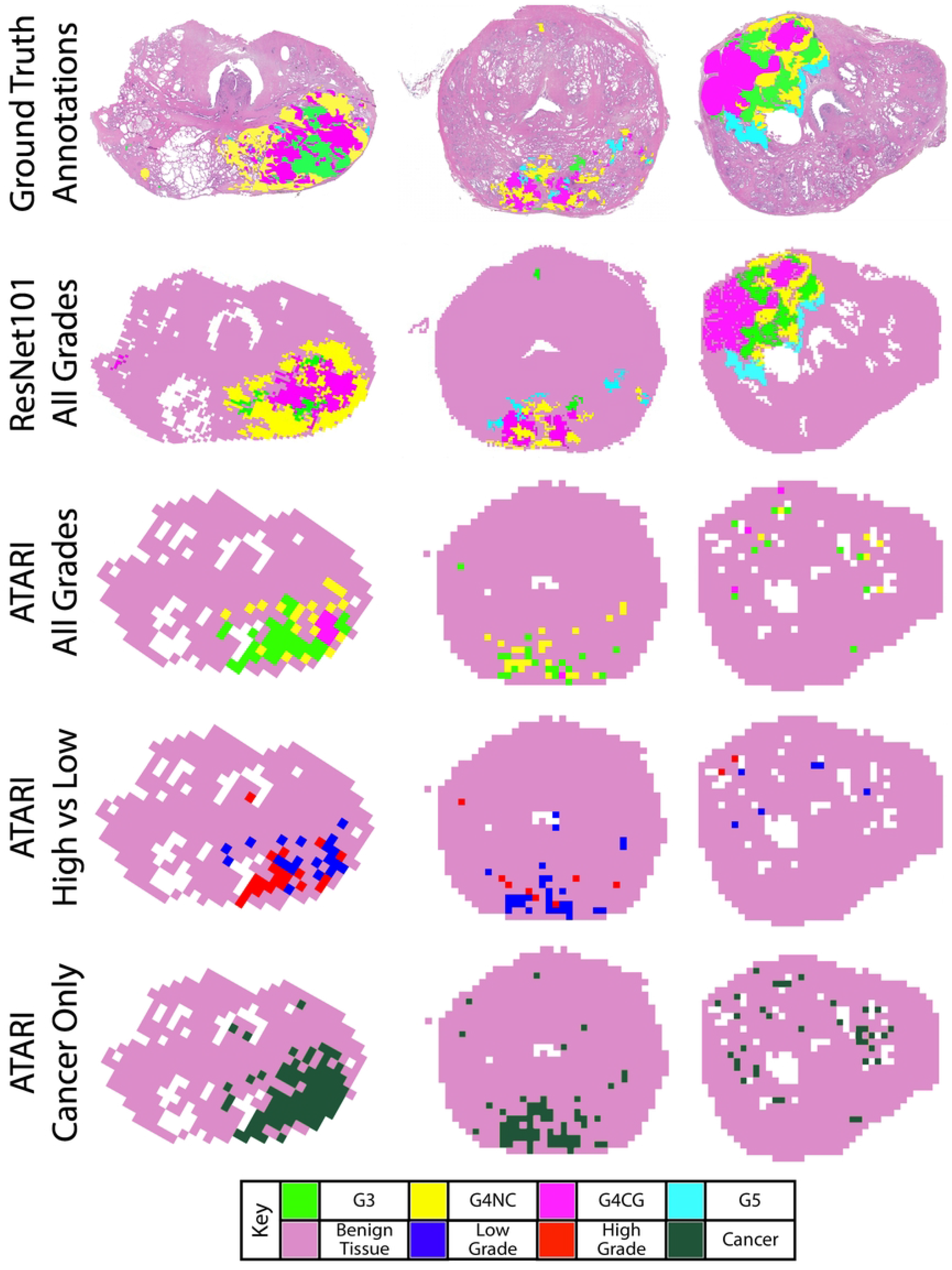
Ground truth annotation maps compared to the ResNet101 model for all Gleason grades and the three tested ATARI models: all Gleason grades, high-vs low-grade cancer, and cancer vs non-cancer only. ResNet101 model for all Gleason grades and the three ATARI models: all Gleason grades, high-vs low-grade cancer, and cancer vs non-cancer only.

## Discussion

In this study, high-resolution tiles taken from annotated regions on whole-mount digital slides after radical prostatectomy were used to train models to support pathologist diagnoses of prostate cancer. Specifically, the ATARI model used quantitative features to classify glandular features, whereas the ResNet101 classifier used deeper textural features of histology. The ATARI was only able to accurately predict cancer and non-cancer, whereas the ResNet101 classifier was able to further predict unique Gleason grades present on the slide. The results from our study indicate that while machine learning models using calculated features may be successful at differentiating tumor from non-tumor, deeper features found using neural networks can further define unique patterns. This may indicate that Gleason patterns exist beyond simple glandular features and may be more readably quantified using textural features. The absolute accuracies of 89% and 83% for the ATARI and ResNet101 models, respectively, show the need for a more general approach to using machine learning for cancer diagnosis.

Machine and deep learning applications are becoming prominent in clinical research. Machine learning focuses on the use of data and algorithms to imitate the way that humans learn. Data used in machine learning applications are human-derived, quantitative metrics that are then analyzed through statistical methods to make classifications or predictions. Deep learning is a sub-field of machine learning that automates the feature extraction without the need for human intervention. It can uncover more nuanced patterns within the data to generate predictions. In this study, our proposed machine-learning model outperformed the ResNet model at classifying cancer from non-cancer; however, the ResNet could classify unique Gleason grades. This may indicate that the features of Gleason grades do not have strong quantitative differences, but rather texture differences that are discernible using a deep learning model. Other prior studies have shown similar results where a trained deep learning model outperformed a simple model trained on handcrafted features[28–30].

Automated Gleason grading applications have been previously applied for multiple purposes. One prior study trained a convolutional neural network (CNN) using WSI-level features constructed from a CNN-based PCa detection model that was trained from slide-level annotations to predict the final patient Gleason Grade Group[31]. This model achieved a 94.7% accuracy at detecting cancer and 77.5% accuracy at predicting the patient Grade Group. While promising, this model does not provide histological annotations to WSI, but rather only predict patient Grade Group. Several previous studies have applied deep learning models to prostate biopsy specimens[11, 32, 33]. While these models have achieved high accuracies at annotating biopsy cores, our ResNet101 model was able to annotate whole-slides images and could distinguish between regions of Gleason 4 cribriform and non-cribriform tumors.

Integrating rapid annotation of Gleason patterns after tissue resection into the clinical workflow could save a tremendous amount of pathologist time. Once slides are digitally scanned, a diagnosis could be predicted automatically based on the automated annotations. This could then be used to rank slides by order of importance for pathologist review and to aid in treatment planning. The proposed models could be applied to large data sets and would decrease the workload on pathologists. Additionally, annotations provided from quantitative metrics may eliminate variability in Gleason annotations.

One major limitation of the study is the use of only one pathologist for annotating the training and test datasets. Inter-observer variability is a known issue in prostate cancer diagnosis, and thus should be addressed in the training phase. Additionally, only one slide scanner was used to digitize the slides used in this study. Future studies should investigate the impact additional slide scanners would have on the generalizability of the models, as this analysis was outside the scope of the current study. Finally, future studies should look at larger populations to provide a more robust dataset of Gleason patterns which may increase accuracy in the machine learning models, as this study had a relatively small cohort of 47 patients.

## Conclusion

We demonstrate in a cohort of 47 patients that machine learning models and neural networks can accurately predict regions of prostate cancer, where the latter network was further able to classify unique Gleason patterns. These models are anticipated to aid in prostate cancer decision support by decreasing the diagnostic burden of pathologists. Future studies should determine how inter-observer and slide scanner resolution impact these networks in their classifications.

## Acknowledgments

We would like to thank our patients for their participation in this study, and the Medical College of Wisconsin Machine Learning Group for helpful feedback and discussion.

## Author Contributions

Conceptualization, S.R.D. and P.S.L.; methodology, A.N., K.A.I. and P.S.L.; software, S.R.D., S.A.B., A.K.L., M.B., A.W., and F.K.; validation, S.R.D., S.A.B., A.K.L., M.B., A.W., F.K., and. P.S.L.; formal analysis, S.R.D.; investigation, P.S.L.; resources, P.S.L.; data curation, P.S.L.; writing—original draft preparation, S.R.D.; writing—review and editing, S.R.D., S.A.B., A.K.L., M.B., A.W., F.K., K.A.I., A.N., and P.S.L.; visualization, S.R.D.; supervision, P.S.L.; project administration, P.S.L; funding acquisition, P.S.L. All authors have read and agreed to the published version of the manuscript.

## References

1. Siegel RL, Miller KD, Fuchs HE, Jemal A. Cancer Statistics, 2022. CA: A Cancer Journal for Clinicians. 2022;72(1):7–33. doi: 10.3322/caac.21708.

2. Epstein JI, Zelefsky MJ, Sjoberg DD, Nelson JB, Egevad L, Magi-Galluzzi C, et al. A Contemporary Prostate Cancer Grading System: A Validated Alternative to the Gleason Score. European Urology. 2016;69(3). doi: 10.1016/j.eururo.2015.06.046.

3. Loeb S, Bjurlin MA, Nicholson J, Tammela TL, Penson DF, Carter HB, et al. Overdiagnosis and overtreatment of prostate cancer. Eur Urol. 2014;65(6):1046–55. doi: 10.1016/j.eururo.2013.12.062.

4. Madabhushi A. Digital pathology image analysis: opportunities and challenges. Imaging Med. 2009;1(1):7–10. doi: 10.2217/IIM.09.9.

5. Niazi MKK, Parwani AV, Gurcan MN. Digital pathology and artificial intelligence. Lancet Oncol. 2019;20(5):e253–e61. doi: 10.1016/S1470-2045(19)30154-8.

6. McGarry SD, Bukowy JD, Iczkowski KA, Lowman AK, Brehler M, Bobholz S, et al. Radio-pathomic mapping model generated using annotations from five pathologists reliably distinguishes high-grade prostate cancer. Journal of Medical Imaging. 2020;7(05). doi: 10.1117/1.jmi.7.5.054501.

7. Ozkan TA, Eruyar AT, Cebeci OO, Memik O, Ozcan L, Kuskonmaz I. Interobserver variability in Gleason histological grading of prostate cancer. Scandinavian Journal of Urology. 2016;50(6). doi: 10.1080/21681805.2016.1206619.

8. Arvaniti E, Fricker KS, Moret M, Rupp N, Hermanns T, Fankhauser C, et al. Automated Gleason grading of prostate cancer tissue microarrays via deep learning. Sci Rep. 2018;8(1):12054. doi: 10.1038/s41598-018-30535-1.

9. Bulten W, Pinckaers H, van Boven H, Vink R, de Bel T, van Ginneken B, et al. Automated deep-learning system for Gleason grading of prostate cancer using biopsies: a diagnostic study. The Lancet Oncology. 2020;21(2). doi: 10.1016/S1470-2045(19)30739-9.

10. Lokhande A, Bonthu S, Singhal N. Carcino-Net: A Deep Learning Framework for Automated Gleason Grading of Prostate Biopsies. Annu Int Conf IEEE Eng Med Biol Soc. 2020;2020:1380–3. doi: 10.1109/EMBC44109.2020.9176235.

11. Ryu HS, Jin MS, Park JH, Lee S, Cho J, Oh S, et al. Automated Gleason Scoring and Tumor Quantification in Prostate Core Needle Biopsy Images Using Deep Neural Networks and Its Comparison with Pathologist-Based Assessment. Cancers (Basel). 2019;11(12). doi: 10.3390/cancers11121860.

12. Brehler M, Lowman AK, Bobholz SA, Duenweg SR, Kyereme F, Naze C, et al. An automated approach for annotation Gleason patterns in whole-mount prostate cancer histology using deep learning. SPIE. San Diego, California 2022.

13. Menon M, Hemal AK. Vattikuti Institute prostatectomy: A technique of robotic radical prostatectomy: Experience in more than 1000 cases. Journal of Endourology. 2004;18(7). doi: 10.1089/end.2004.18.611.

14. Sood A, Jeong W, Peabody JO, Hemal AK, Menon M. Robot-Assisted Radical Prostatectomy: Inching Toward Gold Standard. Urologic Clinics of North America 2014.

15. Shah V, Pohida T, Turkbey B, Mani H, Merino M, Pinto PA, et al. A method for correlating in vivo prostate magnetic resonance imaging and histopathology using individualized magnetic resonance-based molds. Review of Scientific Instruments. 2009;80(10). doi: 10.1063/1.3242697.

16. Cox RW. AFNI: Software for analysis and visualization of functional magnetic resonance neuroimages. Computers and Biomedical Research. 1996;29(3). doi: 10.1006/cbmr.1996.0014.

17. Hurrell SL, McGarry SD, Kaczmarowski A, Iczkowski KA, Jacobsohn K, Hohenwalter MD, et al. Optimized b-value selection for the discrimination of prostate cancer grades, including the cribriform pattern, using diffusion weighted imaging. Journal of Medical Imaging. 2017;5(01). doi: 10.1117/1.jmi.5.1.011004.

18. McGarry SD, Hurrell SL, Iczkowski KA, Hall W, Kaczmarowski AL, Banerjee A, et al. Radio-pathomic Maps of Epithelium and Lumen Density Predict the Location of High-Grade Prostate Cancer. International Journal of Radiation Oncology Biology Physics. 2018;101(5). doi: 10.1016/j.ijrobp.2018.04.044.

19. McGarry SD, Bukowy JD, Iczkowski KA, Unteriner JG, Duvnjak P, Lowman AK, et al. Gleason probability maps: A radiomics tool for mapping prostate cancer likelihood in mri space. Tomography. 2019;5(1). doi: 10.18383/j.tom.2018.00033.

20. Iczkowski KA, Torkko KC, Kotnis GR, Wilson RS, Huang W, Wheeler TM, et al. Digital quantification of five high-grade prostate cancer patterns, including the cribriform pattern, and their association with adverse outcome. American Journal of Clinical Pathology. 2011;136(1). doi: 10.1309/AJCPZ7WBU9YXSJPE.

21. Iczkowski KA, Paner GP, Van der Kwast T. The New Realization About Cribriform Prostate Cancer. Adv Anat Pathol. 2018;25(1):31–7. doi: 10.1097/PAP.0000000000000168.

22. Kweldam CF, Wildhagen MF, Steyerberg EW, Bangma CH, Van Der Kwast TH, Van Leenders GJLH. Cribriform growth is highly predictive for postoperative metastasis and disease-specific death in Gleason score 7 prostate cancer. Modern Pathology. 2015;28(3). doi: 10.1038/modpathol.2014.116.

23. van der Slot MA, Hollemans E, den Bakker MA, Hoedemaeker R, Kliffen M, Budel LM, et al. Inter-observer variability of cribriform architecture and percent Gleason pattern 4 in prostate cancer: relation to clinical outcome. Virchows Archiv. 2021;478(2). doi: 10.1007/s00428-020-02902-9.

24. Ruifrok AC, Johnston DA. Quantification of histochemical staining by color deconvolution. Anal Quant Cytol Histol. 2001;23(4):291–9.

25. Breiman L. Bagging predictors. Machine Learning. 1996;24.

26. Breiman L. Random Forests. Machine Learning. 2001;45.

27. Paszke A, Lerer A, Killeen T, Antiga L, Yang E, Tejani A, et al. PyTorch: An Imperative Style, High-Performance Deep Learning Library. Advances in Neural Information Processing Systems. 2019;32.

28. Arunachalam HB, Mishra R, Daescu O, Cederberg K, Rakheja D, Sengupta A, et al. Viable and necrotic tumor assessment from whole slide images of osteosarcoma using machine learning and deep-learning models. PLoS One. 2019;14(4):e0210706. doi: 10.1371/journal.pone.0210706.

29. Sharma S, Mehra R. Conventional Machine Learning and Deep Learning Approach for Multi-Classification of Breast Cancer Histopathology Images—a Comparative Insight. Journal of Digital Imaging. 2020;33:632–54. doi: https://doi.org/10.1007/s10278-019-00307-y.

30. Xu J, Luo X, Wang G, Gilmore H, Madabhushi A. A Deep Convolutional Neural Network for segmenting and classifying epithelial and stromal regions in histopathological images. Neurocomputing. 2016;191:214–23. doi: 10.1016/j.neucom.2016.01.034.

31. Mun Y, Paik I, Shin SJ, Kwak TY, Chang H. Yet Another Automated Gleason Grading System (YAAGGS) by weakly supervised deep learning. NPJ Digit Med. 2021;4(1):99. doi: 10.1038/s41746-021-00469-6.

32. Lucas M, Jansen I, Savci-Heijink CD, Meijer SL, de Boer OJ, van Leeuwen TG, et al. Deep learning for automatic Gleason pattern classification for grade group determination of prostate biopsies. Virchows Arch. 2019;475(1):77–83. doi: 10.1007/s00428-019-02577-x.

33. Nagpal K, Foote D, Tan F, Liu Y, Chen PC, Steiner DF, et al. Development and Validation of a Deep Learning Algorithm for Gleason Grading of Prostate Cancer From Biopsy Specimens. JAMA Oncol. 2020;6(9):1372–80. doi: 10.1001/jamaoncol.2020.2485.

